# Bayesian phylodynamics for developmental biology: incorporating age-dependence

**DOI:** 10.1101/2025.08.28.672870

**Authors:** Nicola Mulberry, Julia Pilarski, Jana Dinger, Tanja Stadler

## Abstract

As novel technologies for single-cell lineage tracing emerge, phylogenetic and phylodynamic tools are increasingly being used to study developmental processes. However, traditional phylodynamic methods, which were originally developed to study viral evolution, rely on assumptions that are difficult to justify in developmental contexts. Notably, due to cells dividing after characteristic generation times rather than after exponential waiting times—as assumed by the traditionally used birth-death model—empirical cell lineage trees deviate from birth-death phylogenies. Here, we present a non-trivial extension of the birth–death phylodynamic model that captures this characteristic feature of development. By applying our method to a public dataset of stem cell colonies, we show how previous estimates of the underlying population-dynamic parameters were biased by the choice of a birth-death tree prior. Beyond developmental biology, our framework provides an approach for analyzing systems where classical birth-death assumptions may be violated or where empirical tree shapes are poorly captured by those expected under standard phylodynamic models. Our method is available as a BEAST2 package.

**Significance:** Applying phylodynamic inference methods to data from developmental biology requires reassessment of the foundational assumptions underlying these tools. We show that cell population dynamics can be captured by an age-dependent branching process, as opposed to the widely used birth-death process. We develop computational methodology for efficient phylodynamic inference under this age-dependent model, thus providing a tool for connecting cell population dynamics to lineage trees. Our method is furthermore, to our knowledge, the first performant implementation of an age-dependent phylodynamic likelihood, and may be more generally applicable to systems which are ill-characterized by traditional birth-death models.

## 1 Introduction

Modern tools in molecular biology are producing large-scale datasets that enable researchers to gain quantitative insights into developmental processes. Using single-cell RNA sequencing techniques, genetic material can be recovered at the resolution of individual genomes and on the scale of up to millions of cells per experiment. These sequencing methods are destructive, so that in most cases we cannot directly observe the cell population dynamics. A key challenge is thus to infer the temporal dynamics of a tissue or other cell population from molecular data obtained from a final sample of cells [1, 2]. The complexity and scale of these datasets demands advancements in computational and statistical methods.

The potential for phylogenetic-based analyses in developmental biology has recently been highlighted [1, 3, 4]. A cellular phylogeny is a graphical representation of the lineage relationships between a sample of cells, where each tip represents a single sampled cell, and branches represent ancestral relationships. Such phylogenies can be inferred using either imaging techniques or genetic data [5]. In the latter case, researchers could use naturally arising genetic variation, or sophisticated lineage tracing techniques to engineer genetic constructs with increased phylogenetic information content [5, 6, 7, 8]. Once a phylogeny has been constructed, it serves as a statistical framework on which to analyse experimental data, such as transcriptomic measurements of cell states [9].

The use of phylogenetic trees as objects from which to infer population dynamics is an established field of study known as phylodynamics [10]. For example, on a phylogeny built from pathogen sequences taken from infected hosts, we can relate branching events to transmission events, thereby allowing estimation of key epidemiological quantities [11]. In macroevolution, each branch corresponds to a particular species, which may give rise to additional species or become extinct; such phylogenies can be used to estimate diversification rates through time [12]. Many such phylodynamic methods are based on the birth-death-sampling model [13, 14], in which a phylogeny is used to fit an underlying stochastic demographic process, together with a specified sampling process. In its basic form, this model assumes that the phylogeny was generated by a simple birth-death branching process [15, 16], along with uniform sampling over extant lineages [17]. These models make the convenient assumption that birth (or branching) events in a phylogeny occur according to a Poisson process, which assumes that the lifetimes of individuals being modelled are exponentially distributed, i.e., the process is Markovian. While modern methods have focused on adding complexity to these models by, for example, allowing birth and death rates to vary in time [18], by adding population structure [19], or by incorporating complementary data sources [20, 21, 22], much less attention has been paid to the underlying Markovian assumption.

There are several challenges in applying phylogenetic and phylodynamic tools to developmental biology. First, to reconstruct cell phylogenies from sequencing data, we need to accurately describe the mutation process. This is particularly important when working with lineage tracing data, where standard assumptions about molecular evolution are violated. This problem has been addressed in previous research [23, 24, 25]. Second, to quantify cell population dynamics from phylogenies, we need to accurately model the underlying tree-generating process. While age-dependent branching processes have long-since been used to model cellular dynamics [26], their incorporation into phylogenetic analysis has been lacking.

The most well-resolved lineage tree is that of the nematode *C. elegans*, which shows highly synchronous and near-deterministic development during early embryonic stages [27]. By contrast, trees generated under a birth-death model are highly asynchronous and imbalanced. While not all organisms may be as synchronous and deterministic as *C. elegans*, it does seem clear that cellular trees differ significantly from trees generated under the birth-death model, particularly during early stages of development. In a recent simulation study, we have reported biases when inferring the rates of cell division and death under phylodynamic model misspecification, i.e., when fitting the Markovian birth-death process to synchronous cell divisions [28]. Thus, we seek to develop a phylodynamic approach which can capture a wider range of realistic cell population dynamics.

In this paper, we present a phylodynamic method which generalizes the traditional birth-death approaches. Specifically, we introduce non-Markovian dynamics: each lineage’s branching rate depends on its internal *age*, and not on an external time scale. Age-dependent processes are well studied in the branching process literature, however, they have seen less application in the field of phylodynamics [12, 29, 30]. While the likelihood presented here was first derived in Jones [31], this is, to our knowledge, the first implementation and application of the method. We furthermore present an efficient and scalable approximation to the exact likelihood which allows us to perform inference in a Bayesian setting. We implement the likelihood calculation in BEAST2 [32], enabling Bayesian phylodynamic analysis under the age-dependent branching model. As a proof-of-concept, we apply our package to genetic lineage recordings of mouse embryonic stem cells [33, 34]. As a result, we characterize the cell population-dynamic parameters and find that parameters inferred from incomplete cell phylogenies under age-dependent branching strongly agree with estimates obtained from ground-truth cell population trees, while the estimates differ when using the traditional birth-death model.

## 2 Results

### 2.1 Phylodynamic likelihood with Age-Dependent Branching (ADB)

In our underlying population-dynamic model, we assume that cells live for some amount of time, after which they either die with probability *d* or otherwise divide, leaving behind two daughter cells. Crucially, this branching rate increases with the age of the cell. Upon division, the age of each daughter cell is set to 0, and the process repeats. This is a significant departure from the birth-death model, wherein division and death occur at some specified rate (either constant in time or dependent on an external time scale). At some final time, extant cells are sampled with uniform probability *ρ*; the resulting phylogeny displays the ancestral relationships among this sample of cells. Here, we model cell lifetimes as being Gamma distributed with some shape parameter *k >* 0 and mean lifetime *ℓ ∈* ℝ_+_. For the approximation, we require *k ∈* ℕ, i.e., Erlang distributed lifetimes. One could interpret the shape parameter *k* as, for example, stages or sub-stages in a cell cycle. This model and its effect on tree shapes is illustrated in Figure 1.

**Figure 1.**
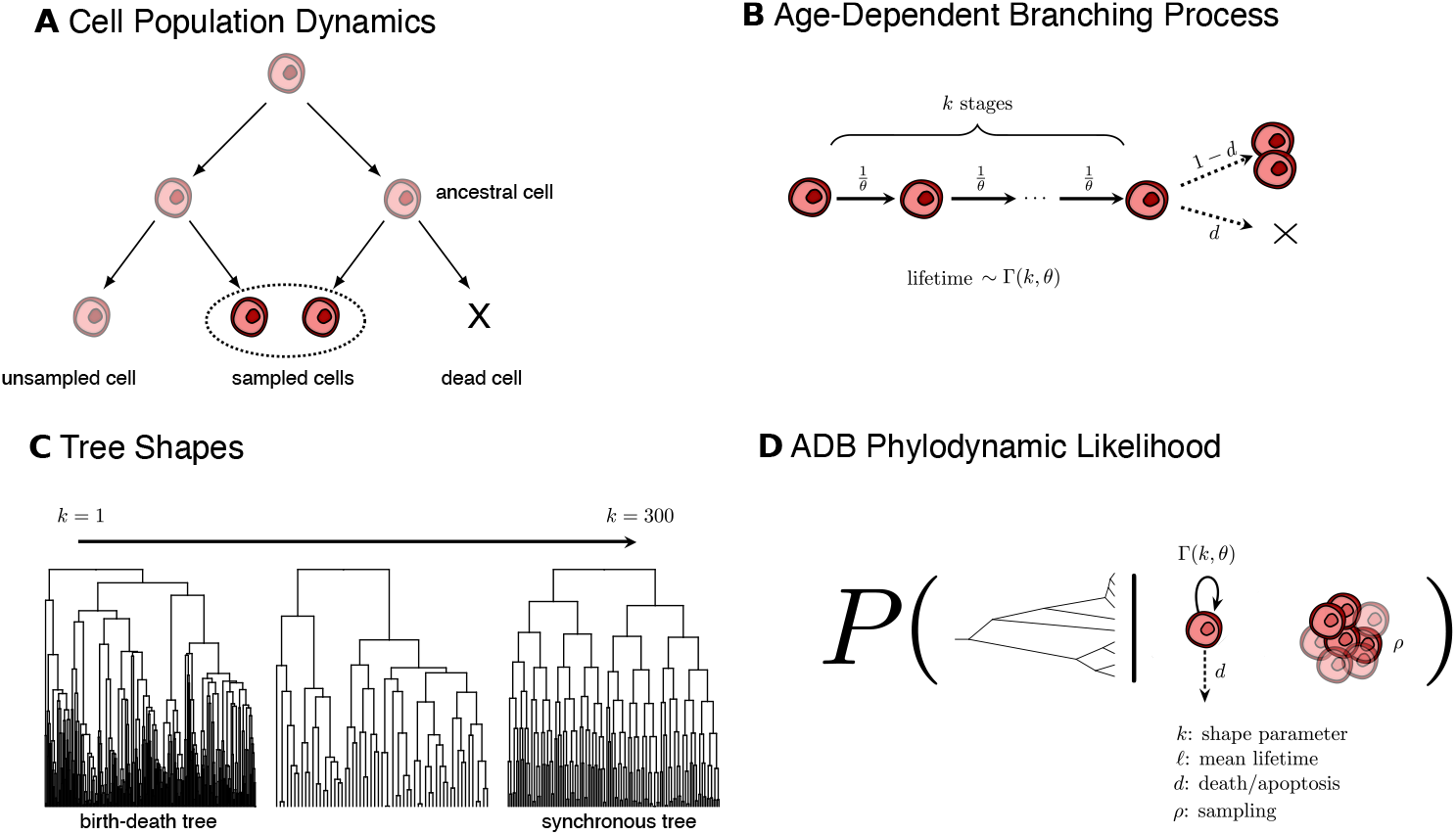
Graphical overview. **A**: Starting from an initial cell, the population undergoes division and death until sampling at a final time point. We only observe a sample of the extant cells, and the cellular history is not directly observed (indicated by shaded cells). **B**: Our method is based on an age-dependent branching process, where we additionally make the assumption that cell lifetimes are Erlang distributed. Solid arrows denote rates, dashed arrows denote probabilities. The birth-death model is a special case of our model found by setting *k* = 1, with corresponding birth rate *λ* = (1 *− d*)*/θ* and death rate *µ* = *d/θ*. **C**: Examples of different tree shapes under various shape parameters *k*, with fixed *d* = 0.01, *ρ* = 0.1 and mean lifetime *ℓ* = 1. As *k* increases, the trees become more and more synchronous. **D**: The phylodynamic likelihood implemented in the BEAST2 package ADB. We parameterize the model by: *k*, the shape parameter of the Erlang distribution; *ℓ*, the mean lifetime (*ℓ* = *kθ*, where *θ* is the scale parameter of the Erlang distribution); *d*, the death probability; and *ρ*, the sampling probability at the final time point.

Section 4.1 presents the derivation for the likelihood of a phylogeny under this model. Note that a lineage is included in the phylogeny only if it has at least one sampled ancestor at the present. While a more detailed derivation is given in Jones [31], we present a condensed version for the single-type case here. Unlike in the Markovian birth-death case, this likelihood has no analytical form, and is instead described by a system of integral equations. In our efficient implementation of the likelihood, we eliminate the need to solve an integral equation over each branch in the tree. Details on our numerical methods as well as our approximations to the exact likelihood are given in Section 4.2.

### 2.2 Tree simulations & lineage-through-time plots

We introduce the R package *scTreeSim* to generate phylogenies under the ADB process (see Section 4.4 for more details). One way to validate these simulations is by comparison to the expected lineage-through-time plots (dLTT), the equation for which is derived in Section 4.3. Overall, we see that the simulator appears to behave as expected (Figure 2).

**Figure 2.**
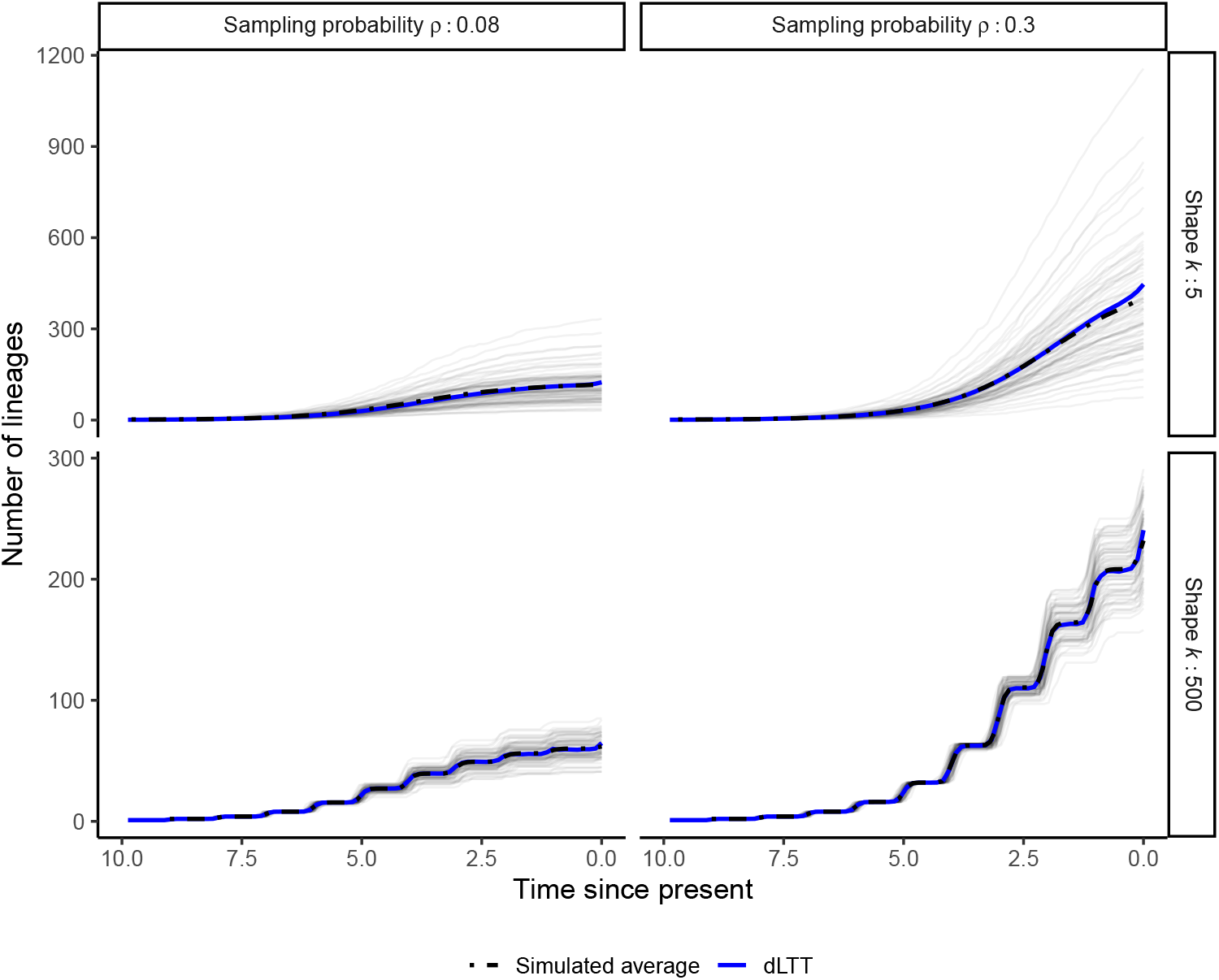
Validation of simulated phylogenies from *scTreeSim* against the expected lineages through time curves (dLTT). Simulations (in grey) are repeated 75 times and the dLTT plot (in blue) is given by Equation (8). The average over the simulated curves is shown in black. The death probability is fixed at *d* = 0 for convenience.

The dLTT curves also give us intuition for our model parameters. We see that, even in low sampling regimes, there is a strong characteristic profile associated to the phylogenies under a high shape parameter, indicative of synchronicity.

### 2.3 Simulation study

Applying Bayesian phylodynamic inference to simulated phylogenies shows that our method estimates all ADB model parameters—shape *k*, mean lifetime *ℓ*, death probability *d*, and sampling probability *ρ*—reliably. The inferred parameters are well-correlated with the true values and satisfy the 95% coverage criterion [35] (Figure 3). Further details are provided in Sections 4.5 and 4.6. In general, mean lifetimes are inferred most accurately, while death probabilities exhibit the largest bias.

**Figure 3.**
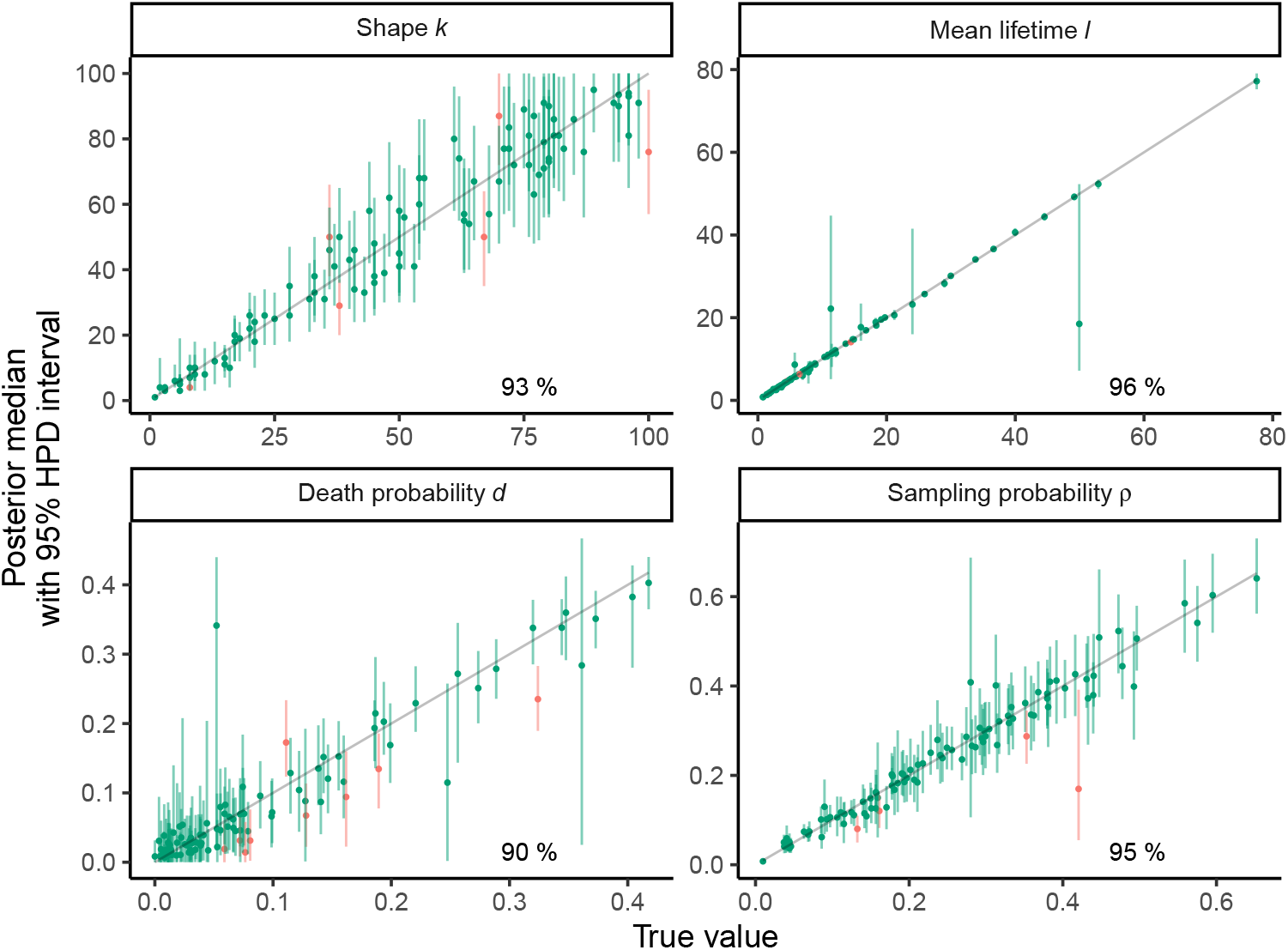
Validation of ADB model on simulations. Panels show the true parameter values (*x*-axis) plotted against the posterior estimates (*y*-axis) for 99 out of 100 simulations for which the MCMC run reached convergence. Dots indicate the medians, bars the 95% HPD intervals, and the diagonal line shows *x* = *y*. Simulations for which the 95% HPD intervals contain the true value are highlighted in green, otherwise in red. The percentage summarizes the coverage (considering all 100 simulations).

It is well known that for the constant-rate birth-death-sampling model, only two of the three parameters *{λ, µ, ρ}* are identifiable [36]. Typically, researchers will fix either the death or the sampling rate and infer the remaining two parameters under the fixed rate. In our case, this simulation study indicates that we recover this non-identifiability as the shape parameter *k* becomes small. However, we observe improved accuracy and precision of *k, ℓ* and *ρ* estimates with increased shape parameter, for which the branching patterns become more regular. Hence, we hypothesize that the model becomes identifiable for large *k* (see SI for more details).

Second, we simulate phylogenetic trees of varying sizes, ranging from 10 to 5000 tips, under a fixed set of parameters. Overall, the posterior distributions of parameters inferred from larger trees are narrower and more tightly centred around the true values (Figure 4A), as expected with an increase in data. However, we observe a small systematic bias in the inference of death probabilities, which becomes more apparent on larger trees. We hypothesize that the bias results from errors in approximating the branch densities (cf. Section 4.2 and SI). This bias is a trade-off for massively reducing the computational cost of the phylodynamic likelihood calculation. For trees with 5000 tips, a single calculation of the full likelihood (which solves an integral equation over each branch in the tree) takes roughly 6s (Figure 4B), meaning an MCMC chain with 1M steps would run for *≈* 70 days. By contrast, our approximation leads to convergence in 10–20h (Figure 4C).

**Figure 4.**
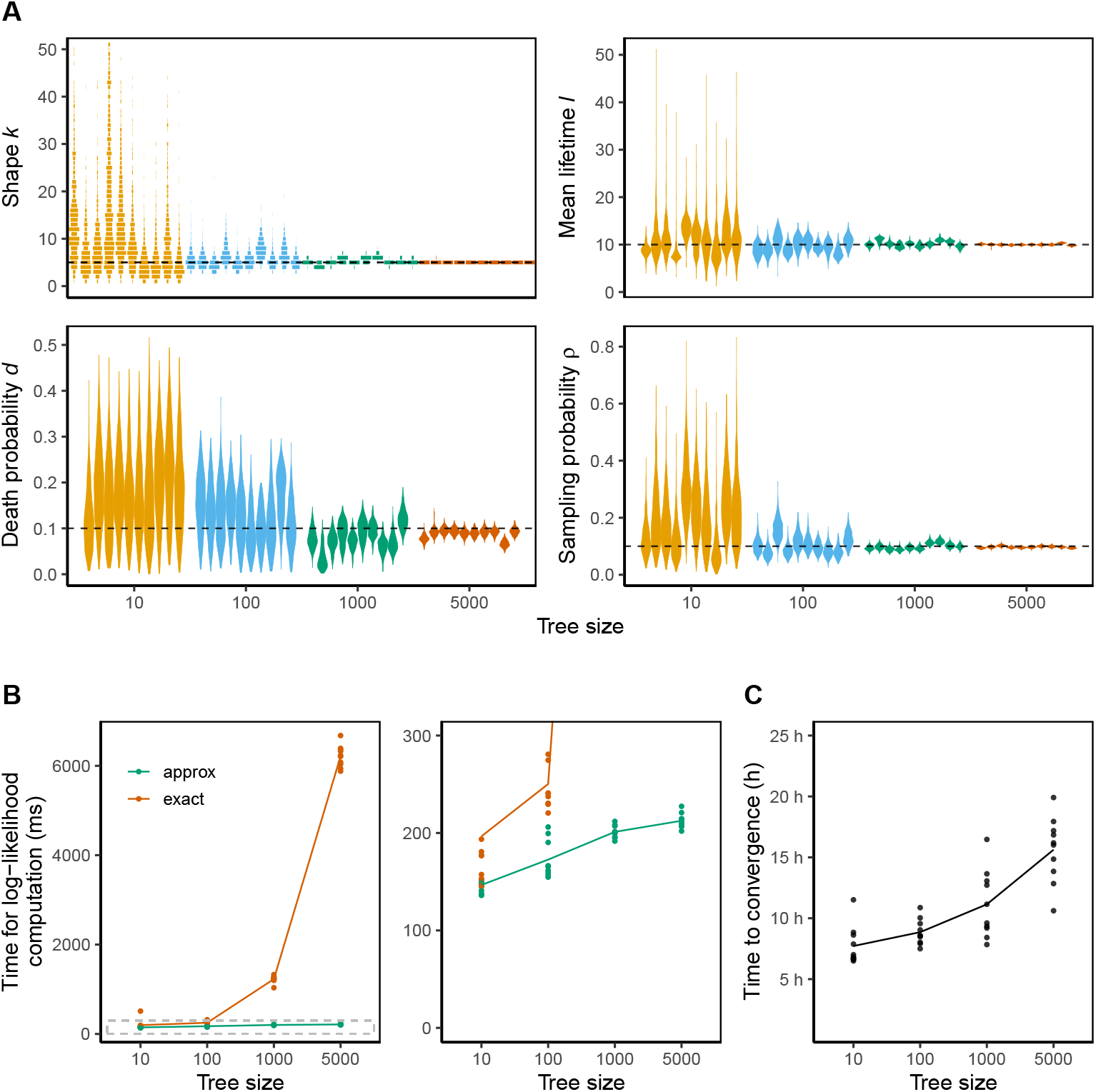
Phylodynamic inference under ADB model on simulated trees of varying size. **A**: Posterior distributions of inferred parameters per simulation, grouped by tree size (i.e., number of tips). **B**: Runtime of phylodynamic log-likelihood computation with exact or approximate edge densities. Dots represents individual trees, lines connect means per tree size. The dashed area in the left panel is magnified on the right. **C**: Inference runtime until MCMC reached convergence threshold (posterior ESS *>* 200). Each dot represents an MCMC chain for a simulated tree.

### 2.4 Application

We demonstrate the ADB model on the experimental *in vitro* dataset of intMEMOIR recordings in mouse embryonic stem cells [33], as available during the DREAM challenge [34]. In the experiment, cells growing into colonies were traced using genetic lineage barcoding and time-lapse microscopy. Crucially, ground-truth cell phylogenies and cell population trees per colony have been extracted from the recordings. These provide us the rare opportunity to evaluate how well a tree-generating model captures the dynamics of the empirical system.

We quantify the population dynamics of stem cells using Bayesian phylodynamic inference (Figure 5A). Under the ADB model, we infer the shape parameter *k* at a posterior median of 32 (95% HPD interval: [30, 35]). This estimate differs substantially from *k* = 1, indicating a higher level of synchronicity and regularity of cell divisions in the population than would be expected under the birth-death (BD) model. Further, we estimate the mean lifetime *ℓ* of cells to be 12.20h ([12.06, 12.32]) and the death probability *d* to be 0.092 ([0.075, 0.112]). These ADB estimates do not overlap with BD estimates, highlighting that phylodynamic parameters are closely linked to tree shape. Moreover, the posterior distributions of *ℓ* and *d* are narrower under the ADB model compared to the BD model, indicating a higher level of confidence in our estimates. Note that we here fix the sampling proportion *ρ* to the true value, as was done in Seidel and Stadler [23].

**Figure 5.**
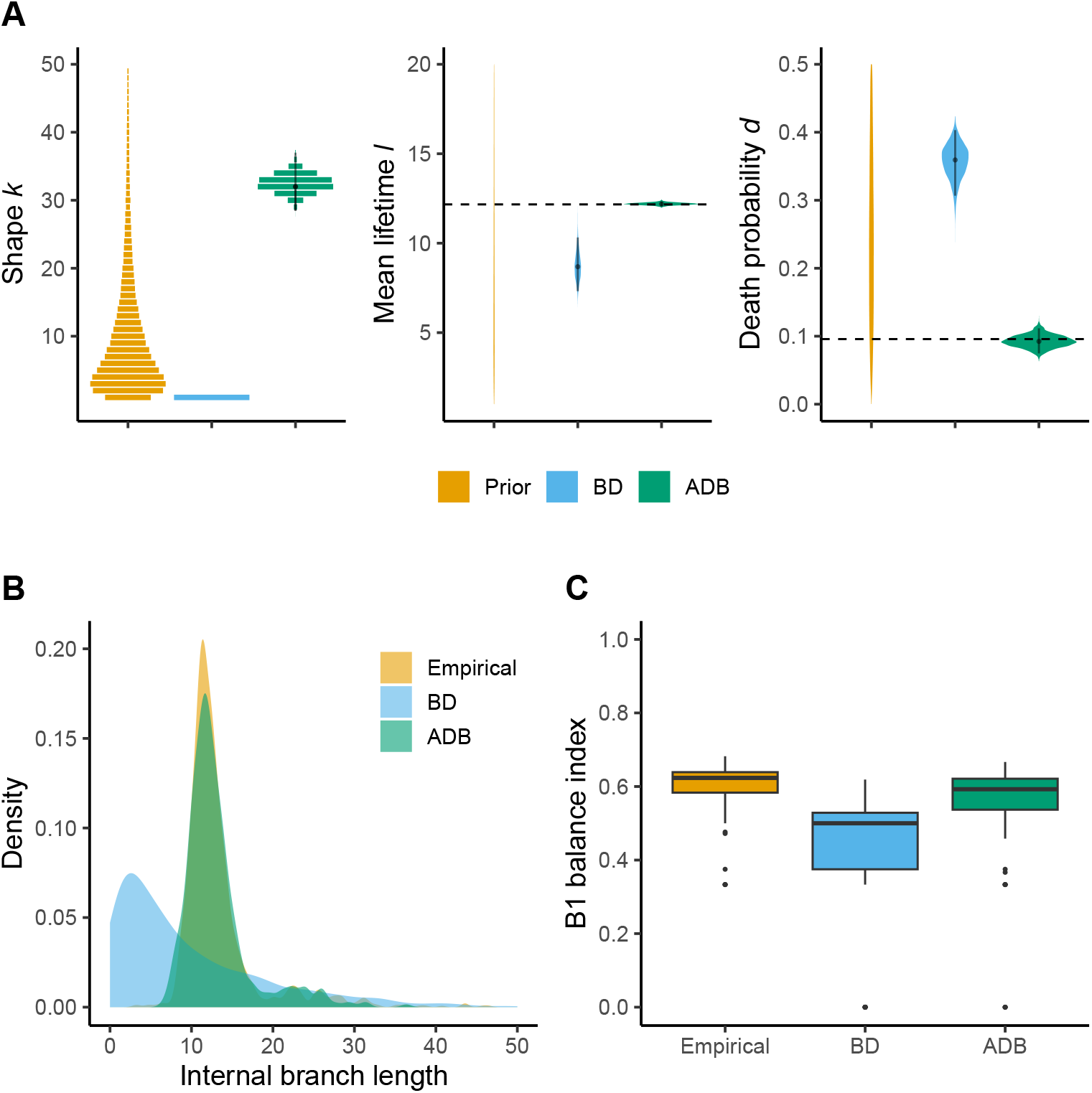
Application of phylodynamic models to single-cell phylogenies from mouse embryonic stem cell colonies. **A**: Prior and posterior distributions of phylodynamic parameters inferred from cell phylogenies under the BD and ADB models. Dashed lines indicate the mean lifetime 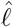 and death probability 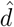 calculated from ground-truth cell population trees (obtained from time-lapse microscopy) for reference. **B**: Internal branch length distribution, and **C**: B1 balance index of empirical (ground-truth) cell phylogenies from the experiment and simulated cell phylogenies.

The inferred parameters are based on phylogenies which delineate the lineage history of only a sample of cells and thus lose information about direct parent-offspring relationships and timings of cell divisions and deaths [4]. To assess whether our proposed phylodynamic model quantifies the cell population process correctly based on the incomplete cell phylogenies, we derive the mean lifetime 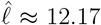 and death probability 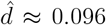 directly from the complete cell population trees. Notably, the empirical estimates fall in the 95% HPD intervals of parameters inferred under the ADB model, but not the BD model. Finally, we compare the empirical cell phylogenies to phylogenies simulated under the inferred BD and ADB models. We observe a strong agreement of the internal branch length distribution and tree balance between empirical phylogenies, and phylogenies simulated under the ADB model (Figure 5B/C).

## 3 Discussion

It is typically not possible to directly observe the full dynamics giving rise to a population of cells; this is particularly true if our only data comes from endpoint single-cell sequencing. However, phylogenetic and phylodynamic tools promise to shed light on these hidden processes. That said, the application of these tools to the developmental context poses numerous challenges on both the phylogenetic and phylodynamic levels. Here, we propose a method to overcome a key challenge on the phylodynamic level: we replace the commonly used Markovian birth-death model with an age-dependent model, taking into account that cell divisions are typically more synchronized than modelled by the traditional birth-death process. Indeed analysis of a single-cell lineage tracing dataset where ground-truth trees obtained through time-lapse microscopy are available highlight that estimates with our novel approach agree with the ground-truth while the classic birth-death approach leads to biased estimates.

Our model assumes that all cells are homogeneous, while typically cell populations are heterogeneous containing cells in different developmental stages. A straightforward extension of our method would be to incorporate multiple cell types. We can model the type changes that occur as cells differentiate, which can furthermore affect growth dynamics of the population, and hence, the tree shape. In this way, phylogenies may provide additional insights into differentiation trajectories. This method would be an extension of the birth-death multi-type (BDMM) method [19, 37].

The current version of the ADB package suffers from a number of limitations. While our implementation enables us to infer phylodynamic parameters in a Bayesian setting on trees of size up to *≈*5000 tips, it nonetheless comes with significant computational costs over the standard, single-type birth-death model (as expected, since this model has an analytical expression for the likelihood). We additionally suffer from significant numerical errors in certain parameter regimes, most notably for very low sampling proportions (less than approx. 1%). This currently poses a problem for applying this method to many developmental data sets which are sparsely sampled. Such regimes may warrant different approaches and approximations, which are left for future work. In addition, our approximation for the edge densities is less accurate under low death probabilities, however we are still able to recover the true parameters in this regime, up to a slight bias in the death parameter on large trees.

We have currently implemented and validated this model as a phylodynamic likelihood; that is, given a fixed phylogeny, we can connect the tree to the underlying population dynamic model. The phylogeny itself is assumed to be either known or inferred from, for example, single-cell lineage tracing data [23, 24, 25, 38]. One can also use the phylodynamic model as a tree prior in a full joint Bayesian inference [32]. However, in our case, joint inference poses additional difficulties since this is a broader tree prior, and the relationship between the shape parameter and tree space has proved challenging to deal with. We hypothesize that new, non-trivial, performant operators are required and leave this for future work.

Identifiability in the context of birth-death phylodynamics is an important concern [39], even when restricting to a class of constant-rate models [36]. In brief, in the constant-rate birth-death case, we can only find two out of the three birth, death, or sampling rates. It is therefore important to investigate identifiability issues under our model, first within the class of constant-rate models. Our initial results, both from simulation studies and from investigating the lineage-through-time plots, indicate that as the true underlying shape parameter increases, we can recover all four population-dynamic parameters (those being, in our case, the shape, lifetime, death and sampling parameters). For shape parameters closer to one, we recover the known constant-rate non-identifiability, at least in the practical sense. A proof for this (non-)identifiability is, however, currently lacking, and will be left for future work.

While the motivation behind this method comes from cellular biology, our method would be applicable to any phylodynamic setting where age-dependence could be a factor, and where birth-death trees do not adequately describe empirical tree shapes. For example, certain species trees have been shown to be less balanced than expected under the birth-death-sampling model [40, 41], a phenomenon which could be captured in our model by taking a shape parameter *less* than one. This would be incompatible with our current approximation, however alternative approaches to address scalability could rely instead on the high sampling rates common in such settings [42].

We have here provided the first, to our knowledge, implementation and application of a non-Markovian generalization of birth-death phylodynamics, and have shown how this method can elucidate cellular population dynamics. Although this method appears promising for analyzing trees in developmental biology, we have nonetheless highlighted remaining challenges that we hope to address with future work.

## 4 Methods

### 4.1 Governing equations

We describe a binary branching process in which the lifetime of a cell is distributed according to a distribution with probability density function *f* (*a*) and cumulative density function *F* (*a*), where *a* denotes the age of the cell. At the end of a cell’s lifetime, it dies with probability *d*, else it divides and generates two daughter cells. We will assume that *d <* 1*/*2. After a fixed amount of time, the extant cells are sampled uniformly with probability *ρ*. Following convention, we track time backwards from the present, with *t* = 0 denoting the present day/sample time and *t* = *t*_*or*_ being the origin time of the process.

We will first derive *P*_0_(*t*), the probability that a cell originating at time *t* will have no descendants in the sample. We will then use this quantity to derive *P*_1_(*t*), the probability that a cell born at time *t* leaves exactly one sampled descendant, and also *b*(*s, t*), the probability that a cell which originated at time *t* had exactly one descendant at time *s*. In this process, there is a fundamental difference between *P*_1_(*t*) and *b*(*s, t*): at the tips a cell’s lifetime is censored due to sampling, whereas along an edge, we observe at least one complete life-cycle.

#### No sampled descendants

Consider a cell which originated at time *t*, where time is counted backwards from the present (*t ≡* 0). There are three possibilities for the cell to have no descendants at present day: either the cell persists for time *t* and then is not sampled at present, it terminates sometime before the present and leads to a death, or the cell divides after some time *w* but its children have no descendants. Therefore, we have

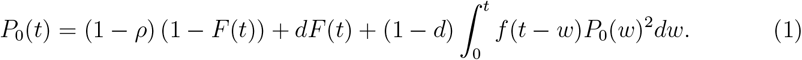

#### Single sampled descendant

A cell originating at time *t* can have exactly one sampled descendant if it survives until present and then is sampled, or if it divides at some time *w ∈* (0, *t*) leading to two daughter cells, one with no sampled descendants and the other with exactly one (but it does not matter which). We get

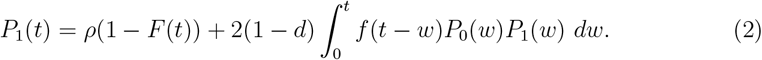

##### Density of an edge (*s, t*)

Consider a cell which originated at time *t*. We wish to work out the probability that this cell has exactly one descendant at time *s >* 0. To compute this quantity, we consider again two possibilities. Either the original cell survives for time *t − s* and then divides, or there is a division at some time *w ∈*(*s, t*) which results in two daughter cells. As before, one daughter cell must leave no sampled descendants. The other daughter cell must result in the edge now between time *s* and *w*. This gives

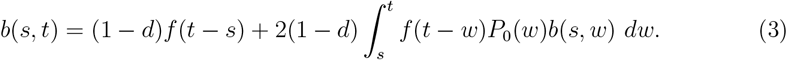

##### Phylodynamic likelihood

Consider a reconstructed tree *𝒯*_*r*_ on *n* tips. Let *t* = 0 denote the present day, and *t* = *t*_*or*_ be the time-of-origin of our process. Let *ε* denote the external edges of *𝒯*_*r*_ and *ℐ* denote the internal edges of *𝒯*_*r*_, and let (*s*_*e*_, *t*_*e*_) denote the start and terminus of edge *e*. The likelihood of a tree is then

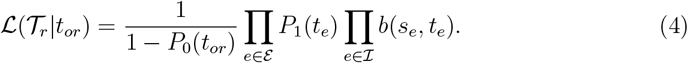

This is essentially the likelihood given in Jones [31], except here we condition on the survival of the process. Considering the likelihood of a labelled tree instead of an unlabelled one, we multiply the equation by factor 2^*n−*1^*/n*! [13].

For example, in the case of a tree with no extinct or unsampled lineages (*ρ* = 1 and *d* = 0), we get

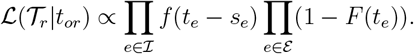

This corresponds to probability of the observed lifetimes, as well as the censored lifetimes at the tips. It is readily verified that Equation (4) simplifies to the analytical expression for the birth-death model with extant sampling [17] when the lifetimes are exponentially distributed.

Alternatively, we can condition on the root of the reconstructed tree, that is, the time of the first cell division event (cf. [17]). Let *t* = *t*_*root*_ and let 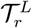 and 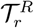 denote the left and right subtree of *𝒯*_*r*_ descending from the root. Then

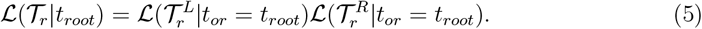

Again, we multiply the equation by factor 2^*n−*1^*/n*!.

### 4.2 Computing the phylodynamic likelihood

Equations (1)–(3) can be solved using fixed point iterations (as suggested also in Jones [31]). For example, to solve for *P*_0_(*t*) (1), we take *X*_0_ = (1 *− ρ*)(1 *− F* (*t*)) + *dF* (*t*) and iterate:

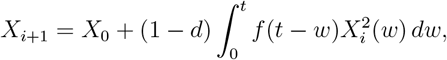

until a convergence criterion is reached. The convolution term can be solved efficiently using the FFT. For the edge probabilities, we consider *b*(*s, t*) to be a function of *t* for each time *s*. Therefore, for each edge *e* = (*s*_*e*_, *t*_*e*_) *∈ ℐ*, we solve *b*(*s*_*e*_, *t*) for *t ∈* [*s*_*e*_, *t*_*e*_], and then evaluate *b*(*s*_*e*_, *t*_*e*_). While each equation can be solved relatively efficiently using the procedure outlined above, since we are required to solve an integral equation over each internal edge, this computation is not scalable to large trees. We instead present an approximate likelihood which is scalable and can be used in a Bayesian inference setting.

#### Approximating edge densities

We now let the cell lifetimes be Erlang distributed. That is, *a ∼* Γ(*k, θ*) with integer shape parameter *k*. Under this assumption, we can approximate the edge densities by an explicit equation. Let 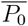 denote the average value of *P*_0_(*τ*) over the edge. Then we let

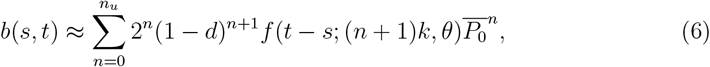

for some finite *n*_*u*_ (see SI for derivation). The number of terms taken in the above approximation depends on a user-specified parameter *ϵ*.

We find that for *k >* 1, solving Eq. (6) is more efficient and scalable than solving the original integral equation for *b*(*s, t*) (3). We note that, under the Erlang assumption, we could instead recast the integral equations as systems of ordinary differential equations by tracking each transitional state explicitly. However, this approach scales poorly with *k* and is not pursued here.

### 4.3 Lineages through Time

Consider a cell that originated at time *τ* in the past. Let *µ*(*t, τ*) be the expected number of lineages at time 0 *≤ t ≤τ* descending from this cell. Following, e.g. Kimmel and Axelrod [26], we get:

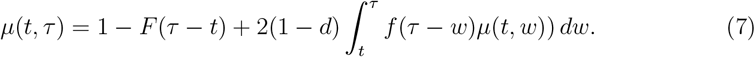

To see this, we first consider that the original cell survives until time *t*, in which case the expected number of lineages stays the same. Otherwise, the cell divides at some time *w ∈* (*t, τ*) with probability *f* (*τ−w*) and survives with probability (1 *− d*), leaving two daughter lineages, each generating *µ*(*t, w*) expected lineages at time *t*. This includes all lineages even if they do not appear in the reconstructed phylogeny, and is conditioned on the time of origin of the process.

In the reconstructed phylogeny, we include a lineage only if it has at least one sampled ancestor at the present. Therefore, the expected number of lineages in *𝒯*_*r*_ becomes

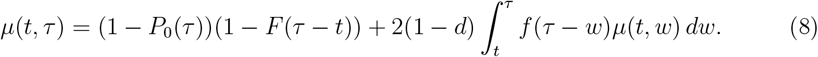

For example, the expected final size is given by *µ*(0, *τ*_*or*_). Typically, we will also condition on the survival of the process.

### 4.4 Tree simulation under ADB

In *scTreeSim*, phylogenetic trees with (1) a predefined number *n* of extant tips or (2) a predefined time of origin *t*_*or*_ can be simulated. We start with a single cell and sample its lifetime from a Gamma distribution with shape *k >* 0 and scale *θ >* 0. At the end of its lifetime, the cell divides with probability (1*−d*) or dies with probability *d*. In the former case, we sample the lifetimes of its two daughter cells. We continue the process until (1) 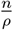 cells are alive at a time, where *ρ* is the sampling probability of extant cells, or (2) the defined time *t*_*or*_ has passed. Next, we set the time to *t* = 0, censor the lifetimes of all living cells, and thus obtain the complete tree. Next, we prune dead cells and sample extant cells uniformly with probability *ρ*. The tree is rooted, binary and ultrametric.

Note that in (1) we simulate the process forward in time until the specified number of living cells is first reached. This procedure, referred to as the simple sampling approach (SSA) can produce biases in the tree origin and the length of external branches [43], as later periods with the same number of living cells are disregarded. In practice, however, these biases only become apparent for higher death probabilities, *d* ≳ 0.4. In this study, we apply our method to lower *d* regimes, hence, we simulate trees with SSA. We provide the general sampling approach (GSA) [43, 44] as an option in *scTreeSim* and recommend using it for higher *d*.

### 4.5 Bayesian inference

Consider a time-scaled phylogenetic tree *𝒯* obtained from empirical observation or computational reconstruction. The goal is to characterize the tree-generating process, that is, the population dynamics which resulted in the observed phylogeny. Using Bayes’ rule, the posterior distribution of phylodynamic parameters is given by

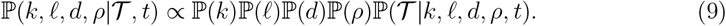

Here, shape *k ≥*1, mean lifetime *ℓ >* 0, death probability *d ∈* [0, 1] and sampling probability *ρ ∈* [0,1] parametrize the ADB process with Erlang-distributed lifetimes. The phylodynamic likelihood ℙ(*𝒯* | *k, ℓ, d, ρ, t*) is given by Equation (4). For inference, prior distributions can be specified for each parameter. In BEAST2, the posterior distribution is approximated by efficiently sampling from the parameter space with the Markov chain Monte Carlo (MCMC) algorithm.

### 4.6 Simulation study

First, we simulate 100 phylogenetic trees with 100 tips under the ADB process using *scTreeSim*. For each simulation, we draw parameters from the prior distributions

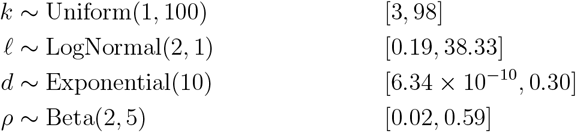

with 95% highest density intervals denoted in brackets.

We then infer *k, ℓ, d* and *ρ* from the trees (with known time of origin) using the BEAST2 package ADB. For each tree, we run an MCMC chain of length 1M for 24h, or until effective sample size (ESS) is 200. We use a sampling frequency of 1000 and discard a 10% burn-in. For each parameter, we summarize the posterior distribution by calculating the median and 95% highest posterior density (HPD) interval.

Note that for one out of 100 trees, the inference did not reach convergence. In this case, the MCMC chain entered an erroneous parameter regime (*d* approaching 0.5 and *ρ* ≲ 0.001, as discussed in SI, Section B), leading to numerical instability in the likelihood calculation and unreliable parameter estimation. These issues are clearly visible in the MCMC traces. Thus, we discard this simulation from further analysis.

To evaluate the inference, we compute the relative bias (i.e., 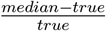), error (i.e., 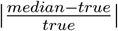), and HPD width (i.e., 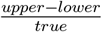, considering the limits of the 95% HPD intervals) of each estimate. We assess the 95% coverage as the percentage of simulations, in which the true parameter value is in the 95% HPD interval of the estimate. For each parameter, we ascertain that the number of 95% HPD intervals containing the true value falls within the expected range (90 to 99, given 100 simulations) [35].

Second, we simulate phylogenetic trees with varying tree size, i.e. 10, 100, 1000, and 5000 tips. We use a fixed set of parameters, i.e. *k* = 5, *ℓ* = 10, *d* = 0.1, *ρ* = 0.1, and generate 10 trees per size. For the inference, we set prior distributions on the parameters as follows:

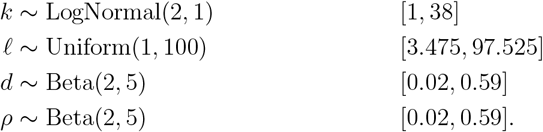

We use the same MCMC settings and summarize the posterior distributions of parameters as before.

### 4.7 Empirical data analysis

We analyze the intMEMOIR dataset [33, 34] as processed by Seidel and Stadler [23]. In the experiment, the development of 106 cell colonies was traced for 54h using time-lapse microscopy and an integrase-based synthetic barcode system. Crucially, some cells in the colonies did not survive until the end of the experiment, dropped out from the recordings, or the readout of their lineage barcode is missing. Thus, the dataset consists of 106 cell phylogenies with 3–39 tips, comprising a sample of extant cells (25 *−* 100%) compared to the recorded cell population trees.

The intMEMOIR recordings have previously been analyzed with the Bayesian phylogenetic framework TiDeTree [23] using the constant-rate BD model with sampling, as implemented in BDSKY [18]. There, the goal was to reconstruct cell phylogenies from sequence alignments by modelling the editing process. In this analysis, we focus on characterizing cell population dynamics based on phylogenies.

For the purpose of a fair comparison, we re-estimate the cell division rate *λ* and death rate *µ* under the constant-rate BD model with sampling, providing ground-truth phylogenies as input to the inference. We condition on the root (i.e. time of the first cell division) and use weakly informative priors on the mean cell lifetime 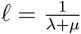 and death probability 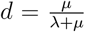. Next, we estimate the phylodynamic parameters *k, ℓ* and *d* under the age-dependent branching model using our package ADB with analogous settings.

For inference, we set the prior distributions as follows:

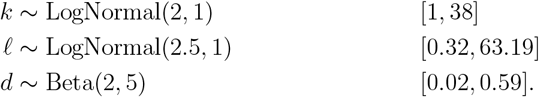

For both phylodynamic models, we run 5 independent MCMC chains with 10^6^ iterations for 24h and verify their convergence. We remove 10% of burn-in and combine the chains to obtain the posterior distribution of parameters.

Independently, we estimate the mean cell lifetime 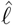 and death probability 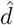 from complete cell population trees. Note that the time scale of the trees is given in movie frames, which were captured approximately every 15 minutes. When converting to hours, the timing of a cell division or death event can only be determined within a 15 minute window around the truth. We derive 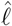 by averaging over the lengths of internal branches and branches ending with a death/ dropout event, and 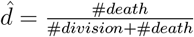 by counting the number of cell divisions and deaths/ dropouts across all trees. We use these empirical measures as reference to assess the accuracy of the phylodynamic parameters inferred from phylogenies.

Next, we simulate 106 phylogenies, each representing the growth of a stem cell colony, under the BD and ADB models. We set the sampling probability per colony to the observed value, and the remaining phylodynamic parameters to the inferred posterior medians. Then, we compute the distribution of internal branch lengths across phylogenies. Additionally, we calculate the B1 index [45] for each phylogeny, normalizing by the number of tips, to compare the balance of empirical and simulated trees.

## 5 Code and data availability

Code to reproduce the analyses presented in this paper is available at: https://github.com/pilarskj/ADB-analysis.

The BEAST2 package is available at: https://github.com/pilarskj/ADB.

Our R package for simulating trees under ADB (along with other single-cell simulation tools) is available at: https://github.com/scCEVO/scTreeSim.

## 6 Author contributions

**Nicola Mulberry**: Conceptualization, Formal analysis, Investigation, Methodology, Supervision, Visualization, Writing – original draft, Writing – review & editing. **Julia Pilarski**: Conceptualization, Formal analysis, Investigation, Methodology, Software, Validation, Visualization, Writing – original draft, Writing – review & editing. **Jana Dinger**: Formal analysis, Investigation, Writing – review & editing. **Tanja Stadler**: Conceptualization, Funding Acquisition, Resources, Supervision, Writing – review & editing.

## 7 Acknowledgements

The authors thank Antoine Zwaans for helpful discussions on the manuscript, and Ugnė Stolz for a code review of the BEAST2 package.

