## supplement for "Bayesian phylodynamics for developmental biology: incorporating age-dependence"

#### A. Approximating edge densities

We now let the cell lifetimes be Erlang distributed. That is, $a \sim \Gamma(k, \theta)$ with integer shape parameter $k$. We first re-write the quantity $b(s, t)$ in an explicit form: the edge $(s, t)$ represents either a single full life-cycle ending in a branching event at time $s$, or two life-cycles with branching events at $w_1 \in (s, t)$ and $s$, and so on. Let $b_n(s, t)$ be the probability that edge $(s, t)$ resulted from $n$ unobserved internal branching events, ending with one at time $s$. Therefore, we let

$$b(s,t) = \sum_{n=0}^{\infty} b_n(s,t), \tag{1}$$

where

$$\begin{aligned}
b_0(s,t) &= (1-d) f(t-s; k, \theta),\\
b_1(s,t) &= 2(1-d)^2 \int_s^t f(t-w_1; k,\theta) f(w_1 - s; k, \theta) P_0(w_1)\, dw_1,\\
&\vdots\\
b_n(s,t) &= 2^n (1-d)^{n+1} \int_s^t \int_s^{w_1} \cdots \int_s^{w_{n-1}} f(w_n - s) \prod_{i=1}^{n} f(w_{i-1} - w_i) P_0(w_i)\, dw_1 \cdots dw_n.
\end{aligned}$$

In the above equation, we take $w_0 = t$ and $f$ to be the pdf of the Erlang distribution.

We now linearize each term by setting $P_0(\tau) \equiv \overline{P_0}$ for $s \leq \tau \leq t$. Under the assumption that the lifetimes are Erlang, we can then write

$$\widetilde{b}_n(s,t) = 2^n(1-d)^{n+1} f(t-s;(n+1)k,\theta)\overline{P_0}^n. \tag{2}$$

We now need to ensure that we take sufficient terms in this approximation. Note that

$$\widetilde{b}_n(s,t) \leq 2^n f(t-s;(n+1)k,\theta),$$

and that the maximum of the equation above occurs around $n^* = \lfloor 2^{1/k}(t-s)/(k\theta) \rfloor$ (in reality, the maximizer of $\widetilde{b}_n(s,t)$ is often lower). Therefore, for some user-specified tolerance $\epsilon << 1$, our approximation thus becomes

$$b(s,t) \approx \sum_{n=0}^{n_u} \widetilde{b}_n(s,t), \tag{3}$$

where $n_u \geq n^*$ such that $\widetilde{b}_n(s,t) \leq \epsilon$.

Under the Erlang assumption, we could instead recast the integral equations as systems of ordinary differential equations by tracking each transitional state explicitly. However, this approach scales poorly with $k$ and is not pursued here. We investigate the error associated with this approximation below.

### B. Accuracy of Likelihood Computation

We first test our implementation of the ADB model in BEAST2 by comparing the likelihood calculation for $k = 1$ to the analytical solution under the constant-rate BD process as implemented in the package BDMM-Prime [1]. We see a good agreement of the likelihood curves with both the "exact" and approximated edge densities (Fig. 1).

However, we observe that in case of very low sampling, i.e. $\rho \lesssim 0.01$, the ADB likelihood deviates from the analytical solution at default settings (Fig. 2). The numerical errors decrease at higher step size for FFT in $P_0$ and $P_1$ calculations. For larger $k$, the errors are further reduced by forcing $P_0$ to be non-increasing if $1-\rho > d/(1-d)$, and non-decreasing otherwise. We find that the low $\rho$ and high $d$ parameter regimes are particularly difficult to resolve, and thus, the numerical solution requires taking more sample points than the default setting.

Conversely, we find that in higher sampling regimes, i.e. $\rho \gtrsim 0.1$, the ADB likelihood is accurate already at a lower than default step size for FFT (Figure 3). Note that reducing the step size substantially reduces the runtime per likelihood calculation.

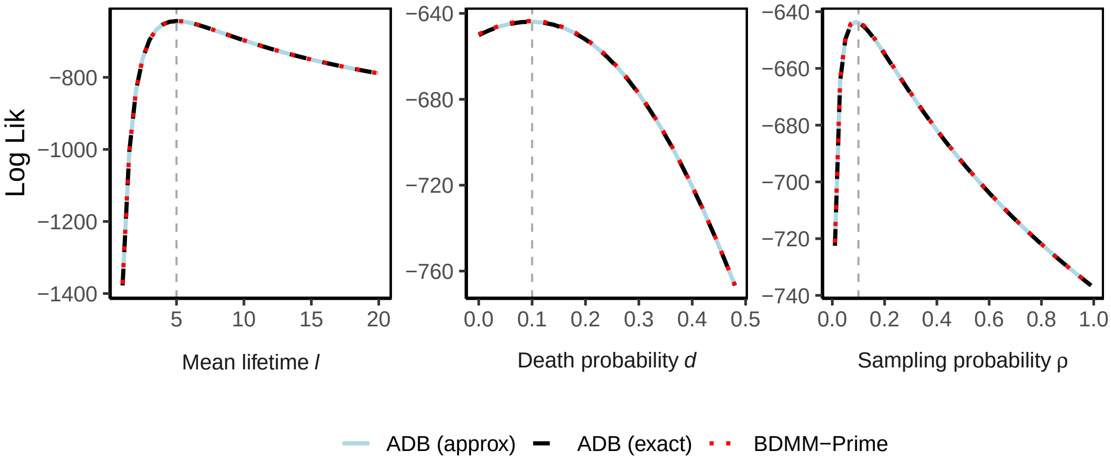


Figure 1: Match of likelihood curves calculated with ADB and BDMM-Prime for $k = 1$. The graphs show the log-likelihood ($y$-axis) for different parameter values ($x$-axis) for a tree with 100 tips, $t_{or} = 50$, and true parameters $\ell = 5, d = 0.1, \rho = 0.1$ (indicated by dashed lines) simulated with TreeSim [2]. The ADB parameters correspond to birth rate $\lambda = 0.18$ and death rate $\mu = 0.02$ in the constant-rate BD process.

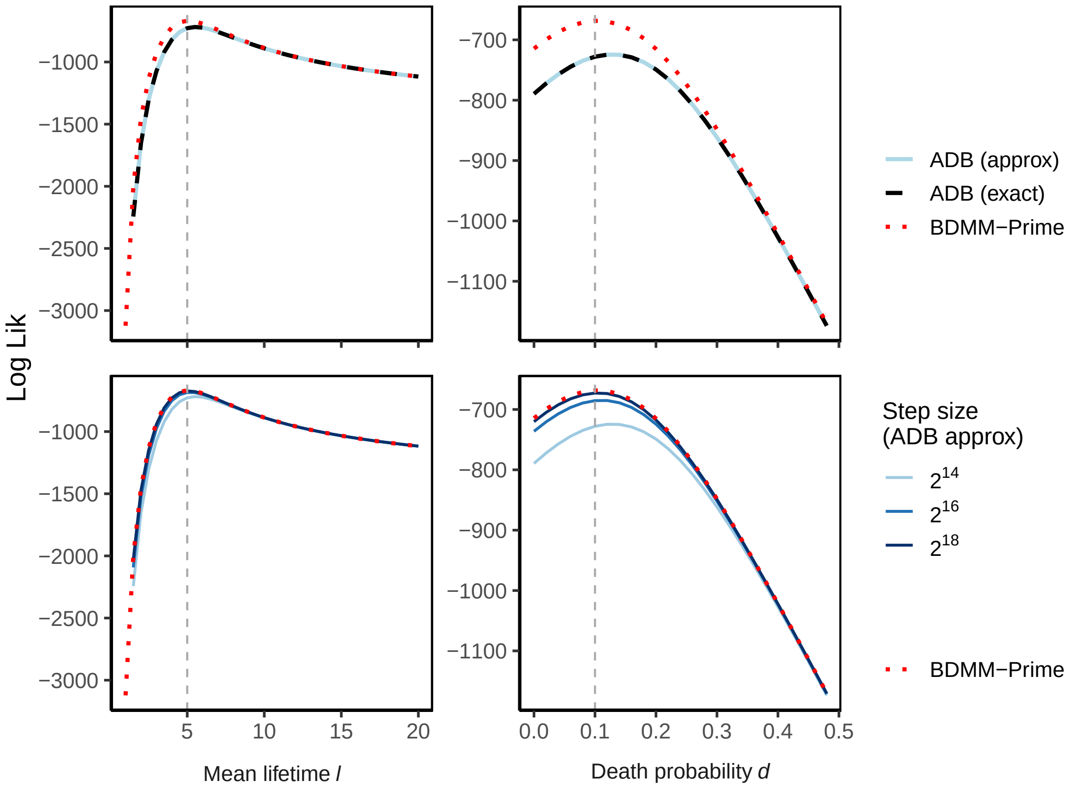


Figure 2: Likelihood curves for $k = 1$ at low sampling, i.e., $\rho = 0.001$, calculated for a tree with 100 tips, $t_{or} = 100$, and true parameters $\ell = 5, d = 0.1$ (indicated by dashed lines) simulated with TreeSim. Comparison of ADB and BDMM-Prime (top). The error decreases with increasing FFT step size when solving the integral equations for $P_0$ and $P_1$ (bottom).

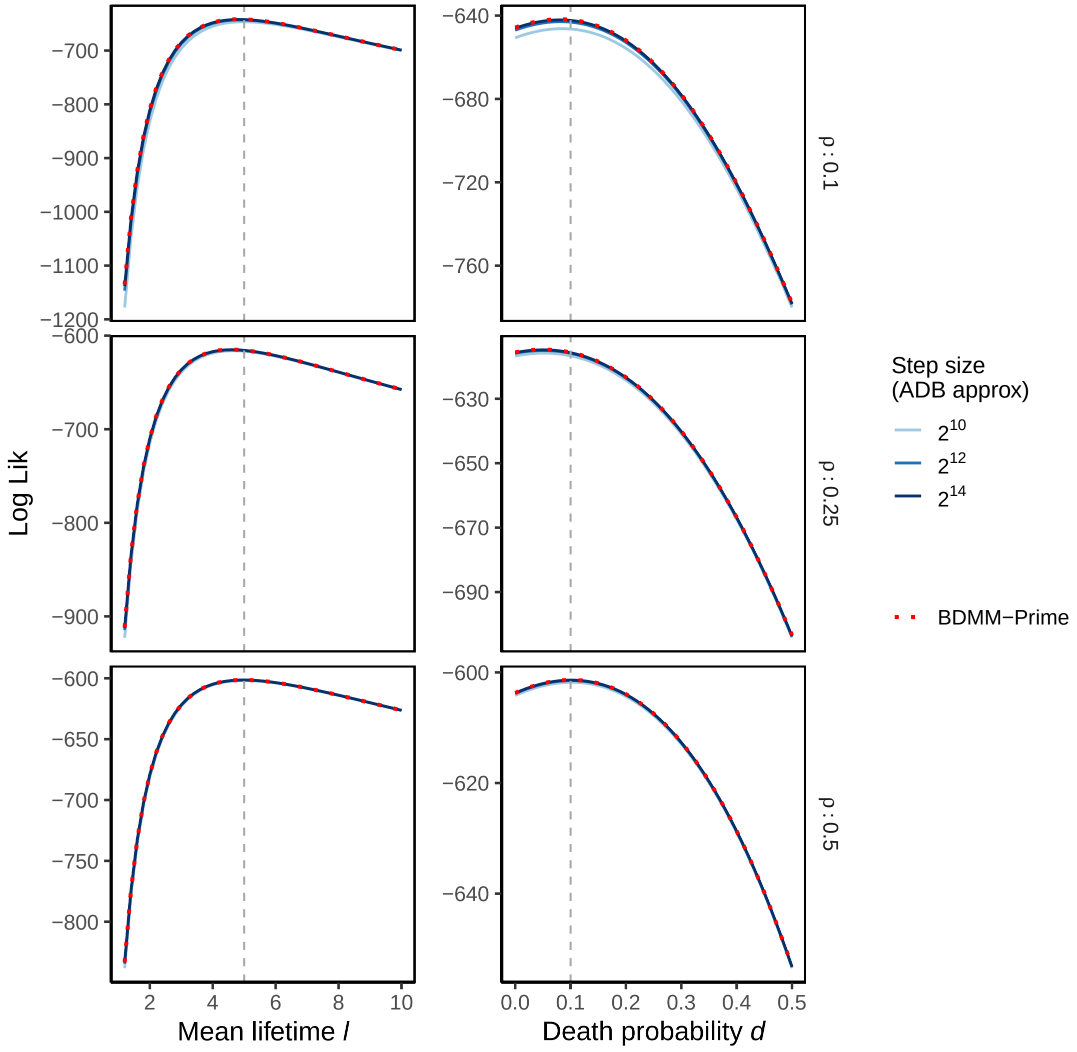


Figure 3: Likelihood curves for $k = 1$ at high sampling, i.e., $\rho \geq 0.1$, calculated for trees with 100 tips and true parameters $\ell = 5, d = 0.1$ (indicated by dashed lines) simulated with *scTreeSim*. Comparison of ADB and BDMM-Prime at varying step size when solving the integral equations for $P_0$ and $P_1$.

Next, we investigate the error in the likelihood calculation due to approximating edge densities (cf. Section 4.2). For each edge $(s,t)$, the error is a combination of the truncation error – which we can control by taking a sufficient number of terms $\tilde{b}_n(s,t)$ to obtain $\tilde{b}(s,t)$ – and an intrinsic error per term $\tilde{b}_n(s,t)$ arising when $P_0$ is not constant along the edge. In this case, $\tilde{b}(s,t)$ might not converge to the exact solution $b(s,t)$. In practice, the intrinsic error results in decreasing accuracy of the likelihood calculation for small $d$, as shown in Figure 4.

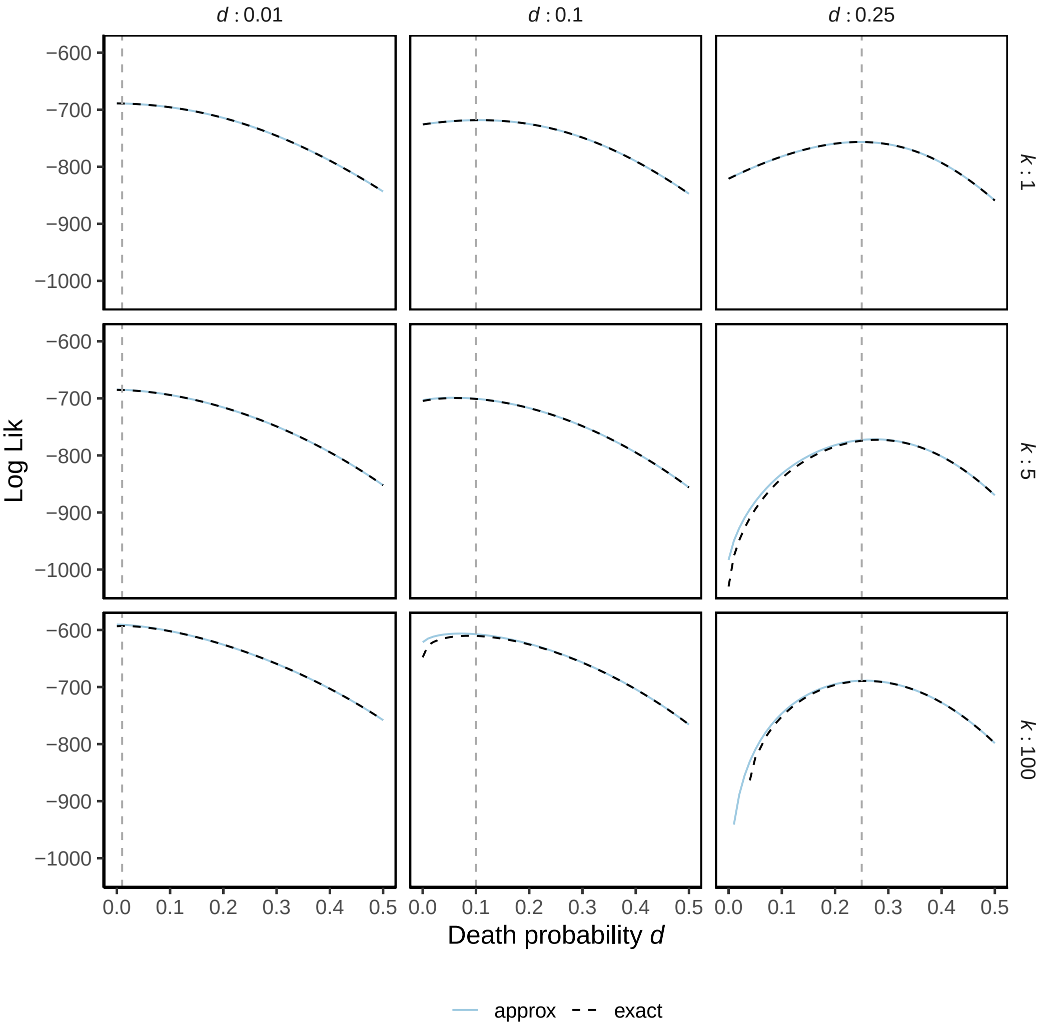


Figure 4: Likelihood curves for trees with varying shape parameter $k$ (rows) and death probability $d$ (columns and dashed vertical lines), $\ell = 10$, $\rho = 0.1$, and 100 tips simulated with *scTreeSim*. The approximation loses precision for small $d$.

### C. Accuracy of Bayesian Phylodynamic Inference

As described in Section 4.6, we simulated 100 phylogenetic trees under the ADB model with parameters drawn from their prior distributions, and re-estimated the parameters using Bayesian phylodynamic inference. Here, we perform coverage validation [3] of the ADB model by calculating the $100 \times \alpha\%$ HPD interval for each tree and parameter, considering a range of credibility levels $\alpha \in (0, 1)$. In Figure 5, we report the percentage of simulations containing the correct data-generating parameter in $100 \times \alpha\%$ HPD interval. We attribute minor deviations from the expected bounds to the relatively low number of simulations, broad priors, and the two sources of errors described before: numerical errors at solving integral equations for $P_0$ and $P_1$, and errors due to approximating the edge densities.

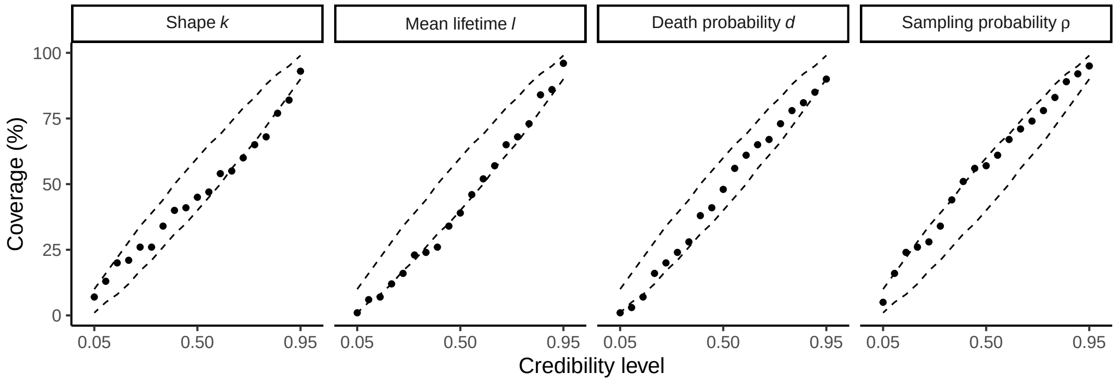


Figure 5: Coverage validation. Dots represent the percentage of simulations ($y$-axis) containing the true parameter in $100 \times \alpha\%$ HPD interval for a range of credibility levels $\alpha$ ($x$-axis). Dashed ellipses indicate expected bounds.

Additionally, in Figure 6, we plot the relative bias, error and 95% HPD interval width of the inferred parameters with respect to true parameter values per simulation.

In our simulation study, we have further observed a systematic bias in the inference of death probabilities on large trees. In Figure 7, we plot the likelihood curves with respect to $d$ for three example trees with 5000 tips. In line with our investigation in Section B, we observe an increasing approximation error for small $d$. This error results in a shift of peaks of the likelihood curves to smaller values. The inferred parameters align with this shift. Note that despite the approximation error, the true parameter $d = 0.1$ has been recovered in 8 out of 10 simulations.

### D. Identifiability

We aim to understand how the known non-identifiability of the constant-rate birth-death-sampling model [4] propagates with higher shape parameters. In particular, assuming the ADB model is the true generative model, we investigate whether all parameters can be estimated. Due to the analytical intractability of our likelihood, we explore here the question of *practical* non-identifiability through simulations and numerical studies.

In our simulation study, we were generally able to infer all four population-dynamic parameters, however we found that parameters inferred from trees with smaller shape parameter $k$ exhibit a larger bias and uncertainty. In this regime, we observe potential parameter non-identifiability, as visible in Figure 8. In this regime ($k \lesssim 10$), while the shape parameters themselves are well-inferred, there is significant uncertainty in the inference of the remaining three parameters. We seek to explore this non-identifiability further.

For the case of shape parameter $k = 1$, we recall the analytical expressions for non-identifiable parameter transformations $(k = 1, \rho, l, d)$ and $(k = 1, \rho', l', d')$ generate equal processes if and only if:

$$l' = \frac{l}{d - (1-d)(1 - 2\rho/\rho')} \quad \text{and} \quad d' = \frac{d - (1-d)(1-\rho/\rho')}{d - (1-d)(1-2\rho/\rho')}. \tag{4}$$

For $k > 1$, we lack an analytical likelihood, and so we investigate possible non-identifiabilities using the deterministic lineage-through-time curves (dLTTs), since any two models with the same dLTT yield identical likelihoods for a given tree [5]. No closed-form solution exists for parameters generating equal dLTTs, but we can draw on numerical methods to generate and compare curves.

We first ask, for a fixed $k$, can we find (practically) congruent processes, one with high sampling and one with low sampling? Figure 9 shows an example where, starting with the known non-identifiability at $k = 1$, we can find qualitatively similar dLTTs in each regime. Finding suitable parameters becomes increasingly difficult as $k$ increases.

In most of our simulation studies, we have found the shape parameter $k$ to be strongly identifiable. An exceptional case occurs when $d$ is very high ($d \gtrsim 0.45$). For large values of $k$, the dLTT approaches a piecewise-constant function with step-size $l$. As a result, different lifetimes can no longer produce identical dLTTs, making theoretical non-identifiability impossible. However, practically speaking, when $d$ is large, we can find parameter sets where a process with a high shape parameter closely approximates one with a lower shape parameter (Figure 10).

All of our simulations thus far have indicated that we can reliably infer all four parameters $(k, d, \ell, \rho)$ as long as we ensure that $d$ is not too large and that we are sufficiently far away from a "birth-death-like" process (i.e. $k >> 1$). In practice, we suggest putting semi-informative priors on the death probability $d$, and to fix the sampling proportion $\rho$ unless there is strong prior information that $k \gtrsim 10$.

### E. intMEMOIR Analysis

As mentioned in Section 4.7, the intMEMOIR dataset contains cell population trees from time-lapse microscopy movies, but also genetic lineage barcodes for a sample of cells per colony [6]. For each tip in a cell phylogeny, i.e., sampled cell at final time point, an array of length 10 indicates the state of the recording DNA sequence at site $i = 1, ..., 10$. Each sequence could either be unedited, or acquire an inversion or deletion – the edits across sites accumulated over time.

Seidel and Stadler [7] have previously analyzed the intMEMOIR recordings with the Bayesian phylogenetic framework TiDeTree. They jointly inferred cell phylogenies and parameters of the editing and phylodynamic models from sequence alignments. Hence, the editing model settings affected phylodynamic estimates. When fixing phylogenies in the inference, the editing and phylodynamic parameters become independent. In this case, the posterior distribution over the space of parameters follows

$$\mathbb{P}(\Phi, \Psi | D, \mathcal{T}) \propto \mathbb{P}(\Phi)\mathbb{P}(\Psi)\mathbb{P}(\mathcal{T}|\Psi)\mathbb{P}(D|\mathcal{T}, \Phi) \tag{5}$$

where $\Psi$ denotes the phylodynamic model parameters and $\Phi$ represents the editing model parameters.

As a consequence, we do not expect the editing model parameters to differ when using different phylodynamic models. Indeed, when jointly inferring editing model parameters with either the BD or ADB model parameters, we obtain consistent estimates, as shown in Figure 11. Furthermore, the estimates agree with the previous analysis of the data, in which the editing rates $r_i$ and the edit-outcome rate multipliers (i.e. for acquiring an inversion $s_i^I$, or a deletion $s_i^D$) are allowed to vary across sites and the following prior distributions are used:

$$\begin{aligned} r_i &\sim \text{LogNormal}(-5, 1) \qquad && [0.0002, 0.035] \\ s_i^I, s_i^D &\sim \text{LogNormal}(0, 1) \qquad && [0.03, 5.17]. \end{aligned}$$

In a supplementary analysis, we run phylodynamic inference on the intMEMOIR dataset with estimating all ADB model parameters, including the sampling probability $\rho$. That is, we infer the shape parameter $k$, mean lifetime $\ell$, and death probability $d$ jointly from all cell phylogenies, and additionally, a sampling probability $\rho$ for each of the 106 trees. We use a broad prior, $\rho \sim \text{Uniform}(0, 1)[0.025, 0.975]$. Despite small tree sizes, and thus noisy input data, we recover the sampling proportion for 84 out of 106 trees. For these trees, the observed proportion is in the 95% HPD interval of the estimate (if $\rho < 1$, otherwise we verify that the posterior distribution approaches 1, specifically, the upper bound of the interval $\geq 0.999$). The mean of the inferred sampling probabilities ($\approx 0.82$) roughly agrees with the mean sampling proportion across trees ($\approx 0.79$). Importantly, the population-dynamic parameters of interest, $k, \ell$ and $d$, could be inferred equally well, when the information on sampling was missing (Figure 12).

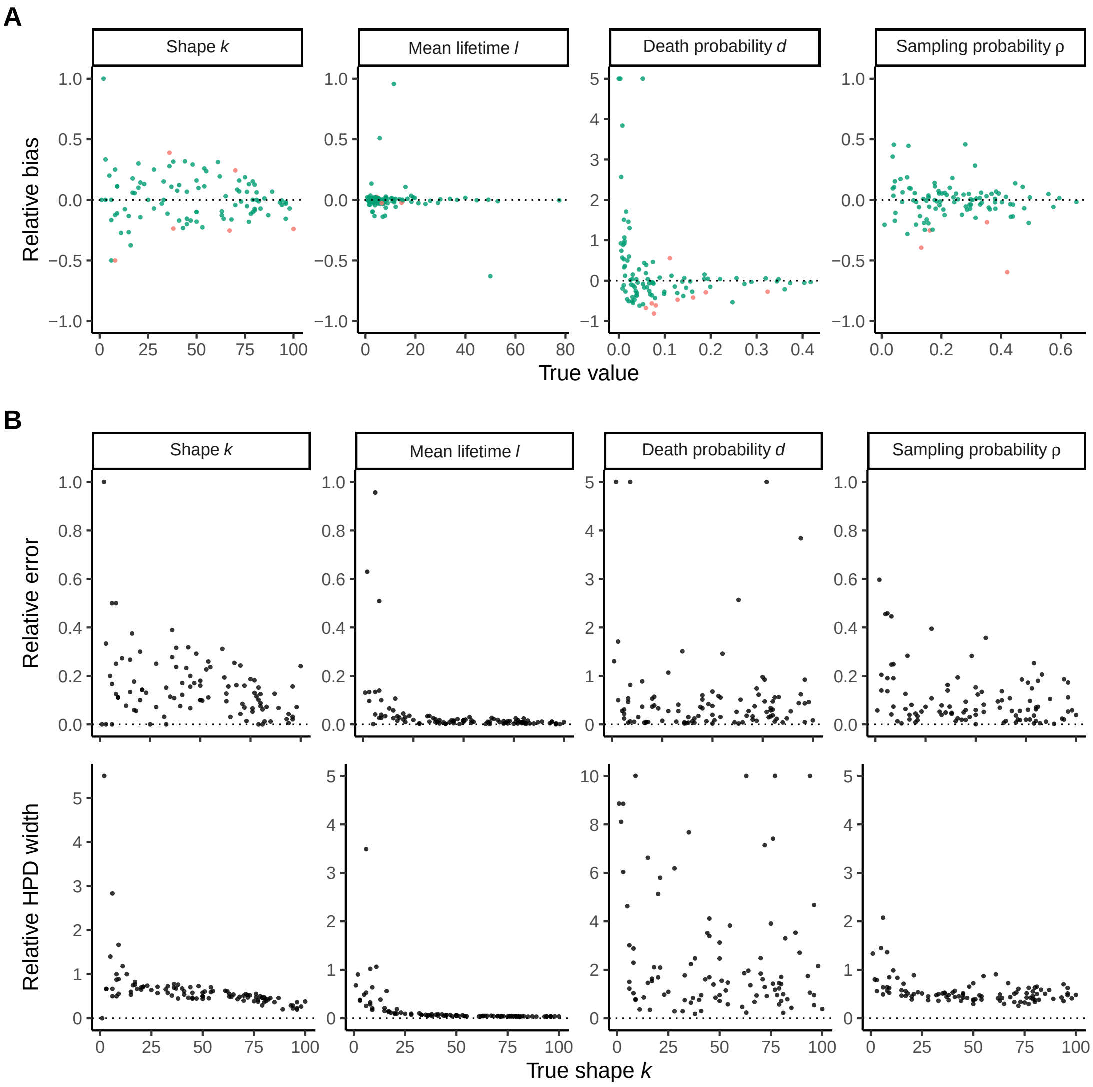


Figure 6: Accuracy of Bayesian phylodynamic inference on simulations. **A**: Relative bias ($y$-axis) of inferred parameters with respect to true parameter values ($x$-axis), colored by 95% HPD coverage. **B**: Relative error (top) and HPD width (bottom) of inferred parameters with respect to true shape parameter $k$. For readability, large values arising through division by near-zero death probabilities are clipped.

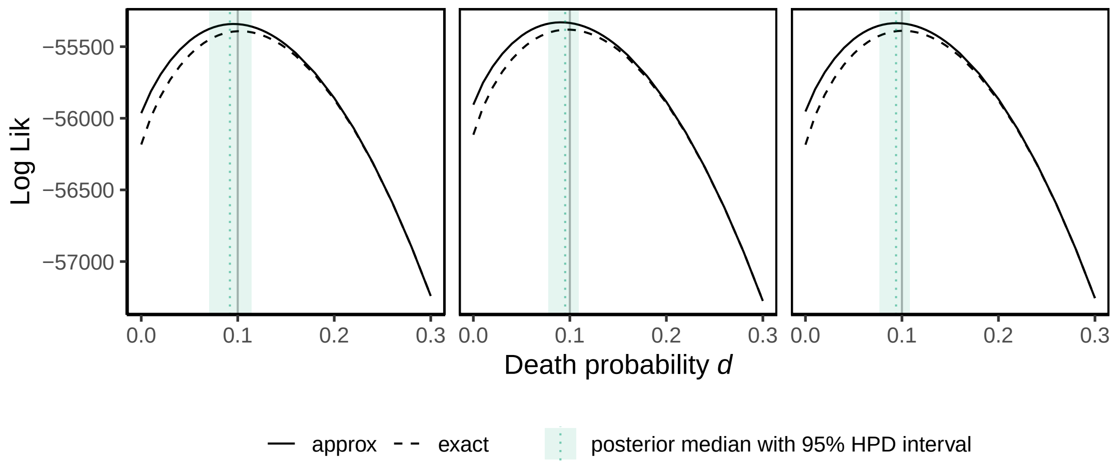


Figure 7: Bias in inferring death probabilities due to approximation error at low $d$. Shown are likelihood curves for three trees with 5000 tips selected from the simulation study. Solid vertical lines indicate the true parameter, dashed lines with ribbon indicate the inferred parameters.

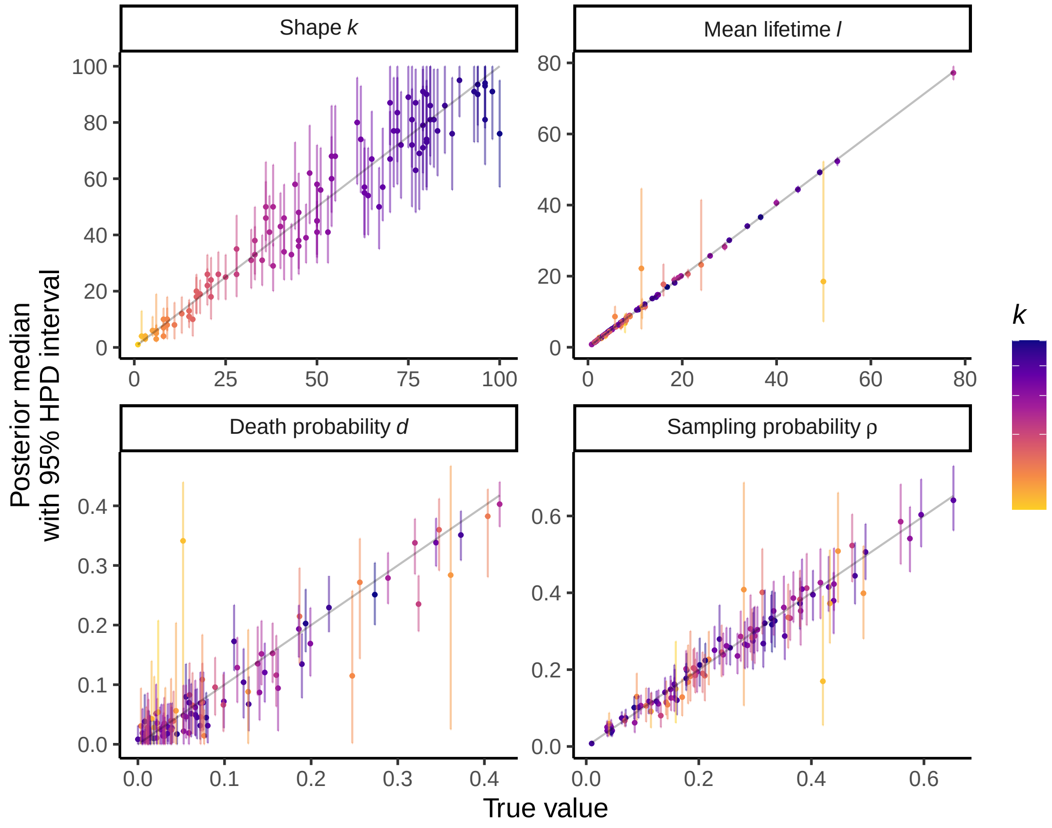


Figure 8: Inference under ADB model at varying $k$. Panels show the true parameter values ($x$-axis) plotted against the posterior estimates ($y$-axis). Dots indicate the medians, bars the 95% HPD intervals, and the diagonal line shows $x = y$. Simulations are coloured by their true shape parameter $k$.

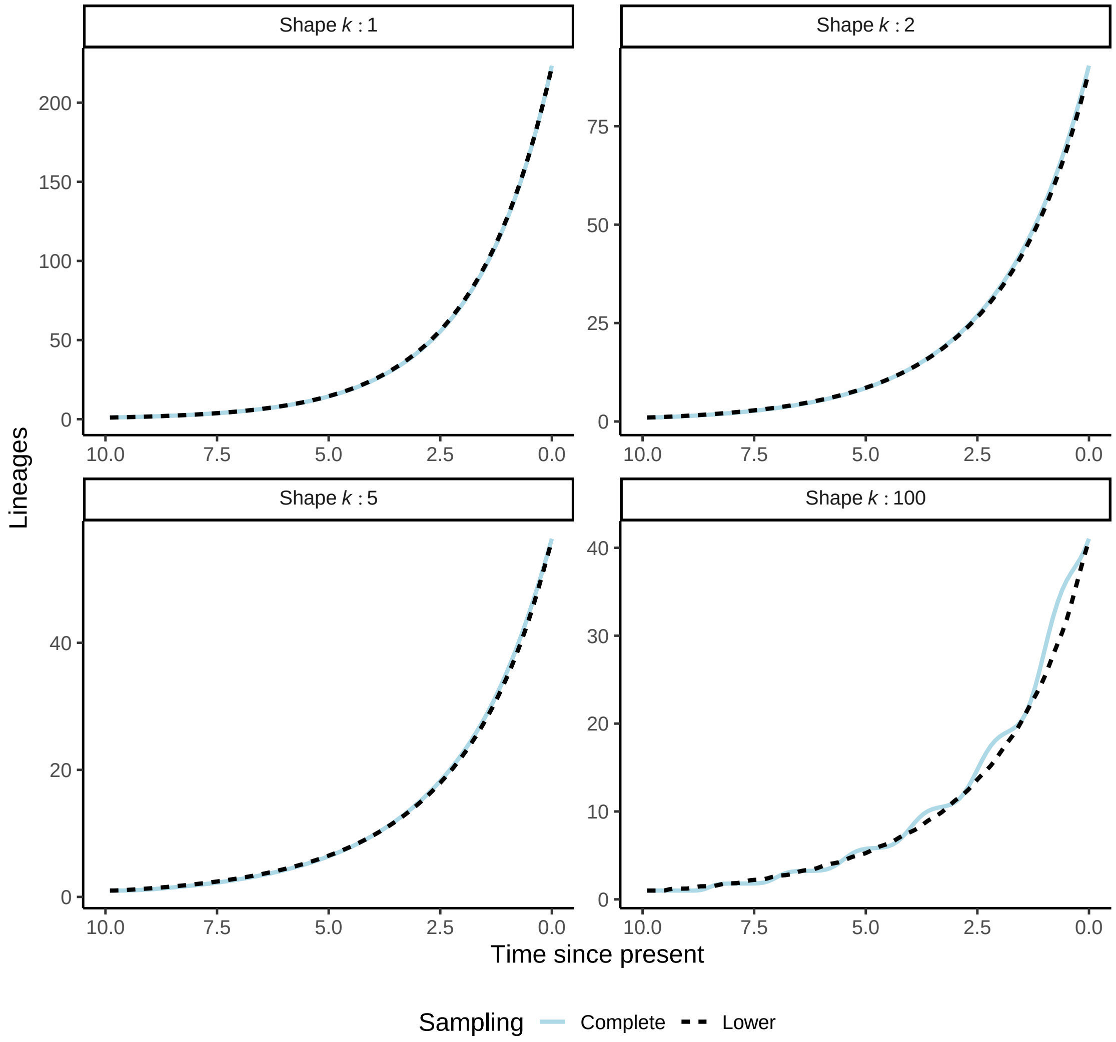


Figure 9: Examples of similar dLTTs. For the fully sampled case, we take parameters $(k, \rho = 1, \ell = 1.5, d = 0.1)$. For each given shape parameter, we set $(k' = k, \rho' = 0.5, \ell' = 0.57, d' = d_k)$, where $d_k$ is the best match found through a grid search.

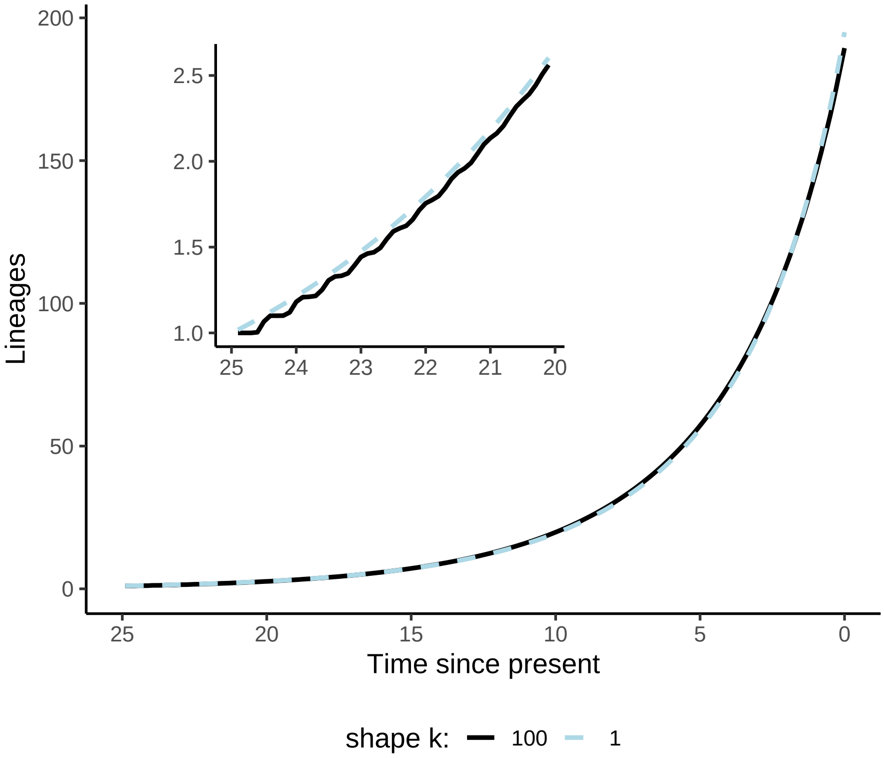


Figure 10: Scenario where a high shape parameter appears to approximate the dLTT with a lower shape parameter. For $k = 100$, we take $d = 0.45$.

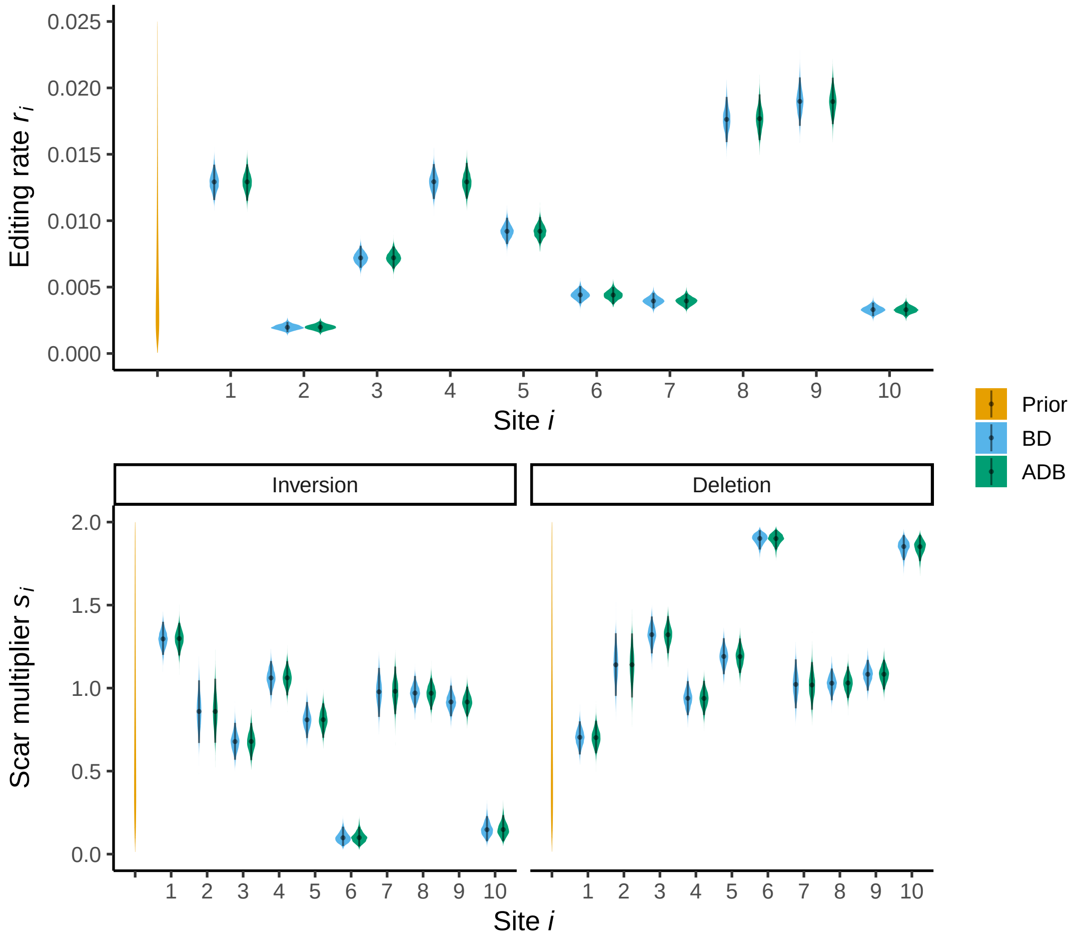


Figure 11: Inference of editing dynamics in intMEMOIR recordings. The graphs show prior and posterior distributions of editing model parameters under TiDeTree, jointly inferred with phylodynamic parameters under the BD and ADB models.

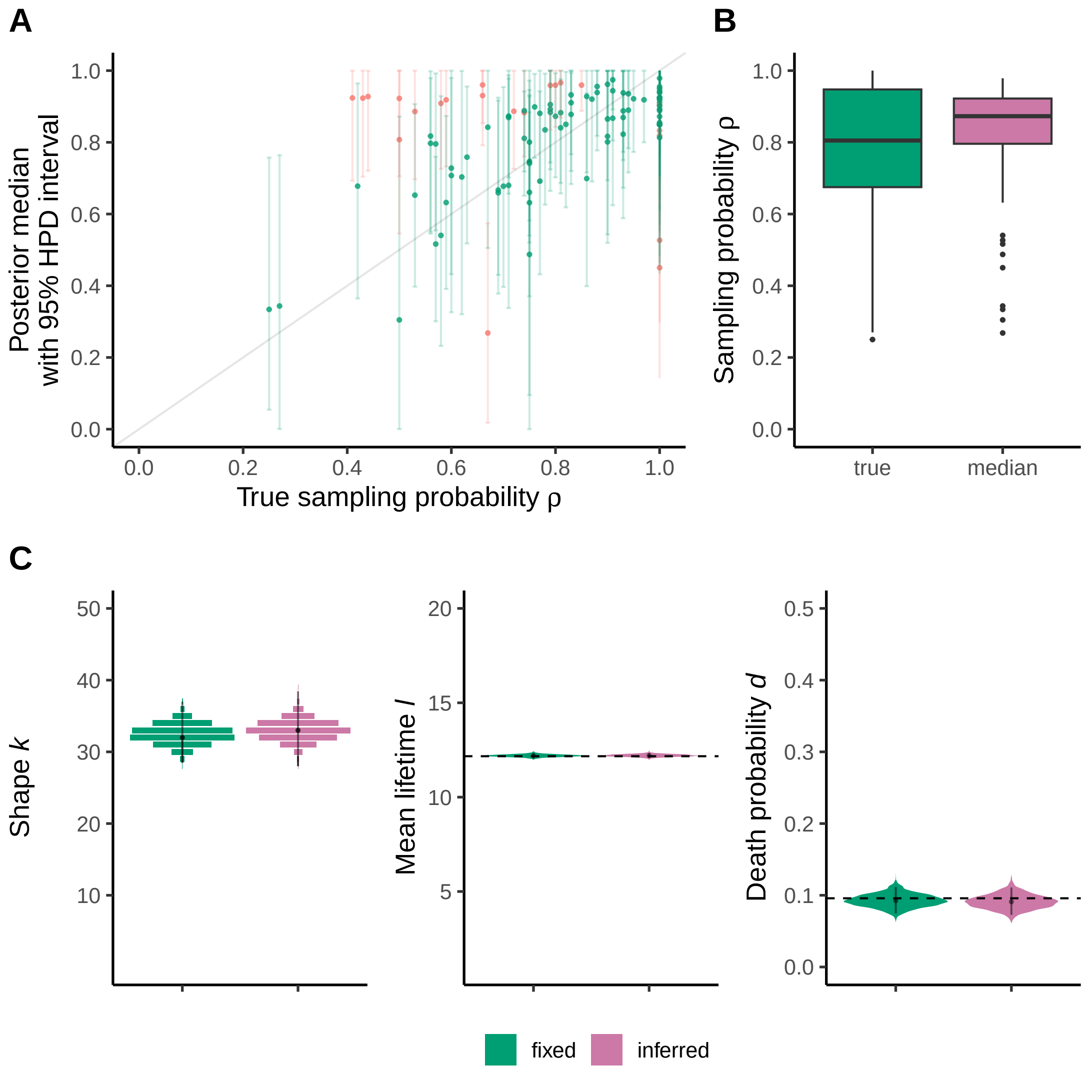


Figure 12: Phylodynamic inference from intMEMOIR cell phylogenies. **A**: Observed sampling proportions ($x$-axis) plotted against the posterior estimates ($y$-axis) per tree. Dots indicate the medians, bars the 95% HPD intervals, and the diagonal line shows $x = y$. Recovered proportions are colored in green, otherwise in red. **B**: Distribution of observed (true) and inferred (median) sampling proportions. **C**: Posterior distributions of phylodynamic parameters inferred from cell phylogenies under the ADB model with fixed or inferred sampling probability $\rho$. Dashed lines indicate the mean lifetime $\hat{\ell}$ and death probability $\hat{d}$ estimated from cell population trees for reference.
